# STOICHIOMETRY OF THE SODIUM PUMP-PHOSPHOLEMMAN REGULATORY COMPLEX

**DOI:** 10.1101/2021.10.12.464104

**Authors:** Jaroslava Seflova, Nima R. Habibi, John Q. Yap, Sean R. Cleary, Xuan Fang, Peter M. Kekenes-Huskey, L. Michel Espinoza-Fonseca, Julie B. Bossuyt, Seth L. Robia

## Abstract

The sodium-potassium ATPase (NKA) establishes ion gradients that facilitate many physiological processes. In the heart, NKA activity is regulated by its interaction with phospholemman (PLM, FXYD1). Here we used a novel fluorescence lifetime-based assay to investigate the structure, stoichiometry, and affinity of the NKA-PLM regulatory complex. We observed concentration dependent association of the subunits of NKA-PLM regulatory complex, with avid association of the alpha subunit with the essential beta subunit followed by lower affinity alpha-alpha and alpha-PLM interactions. The data provide the first evidence that the regulatory complex is composed of two alpha subunits associated with two beta subunits, decorated with two PLM regulatory subunits in intact cells. Docking and molecular dynamics simulations generated a structural model of the complex that is consistent with our experimental observations. We propose that alpha-alpha subunit interactions support conformational coupling of the catalytic subunits, which may enhance NKA turnover rate. These observations provide insight into the pathophysiology of heart failure, wherein low NKA expression may be insufficient to support formation of the complete regulatory complex with stoichiometry (alpha-beta-PLM)_2_.

## Introduction

The sodium pump (NKA, Na^+^/K^+^-ATPase) is an essential membrane-bound transport ATPase establishing vital Na^+^ and K^+^ gradients across the plasma membrane. During its reaction cycle, NKA uses energy from ATP hydrolysis to translocate 3 Na^+^ ions from the cytoplasm into the extracellular space and 2 K^+^ from outside the cell into the cytoplasm. NKA comprises a complex of the α catalytic subunit that binds substrate ions and ATP and a β subunit that is critical for correct localization of the α subunit in the plasma membrane. Na^+^ transport is finely regulated by tissue-specific expression of four different α subunit isoforms with three different β subunit isoforms. Additional homeostatic control provided by interactions of the essential αβ complex with seven diverse regulatory partners from the FXYD family of micropeptides (Geering, 2005; Yap *et al*, 2021). FXYD proteins contain a single transmembrane helix and a highly conserved extracellular extension containing a FXYD motif which gives this family of NKA regulators its name. In the heart, NKA is regulated by an inhibitory interaction with FXYD1, also known as phospholemman (PLM) (Palmer *et al*, 1991). This inhibition results in a comparatively higher intracellular Na^+^ concentration, limiting Ca^2+^ extrusion by the Na^+^/Ca^2+^ exchanger (NCX) and thereby increasing intracellular Ca^2+^. Increased Ca^2+^ handling supports stronger cardiac contractions. This physiological connection between cardiac Na^+^ transport and Ca^2+^ handling provides a mechanism of action for digitalis, an NKA-inhibiting drug used for hundreds of years to increase cardiac contractility in patients with heart failure (The Digitalis Investigational Group, 1997; Newman *et al*, 2008).

X-ray crystallography revealed a NKA regulatory complex with a concise stoichiometry of a single FXYD protein and a single β subunit bound to a single α subunit. However, biochemical studies suggest that the subunits may assemble in a larger functional complex. Initial work by Stein et al. (Stein *et al*, 1973), Ottolenghi and Jensen (Ottolenghi & Jensen, 1983) suggested NKA dimerization based on cooperative ATP hydrolysis, which may indicate a catalytic dimer of functionally coupled α subunits (Askari, 1987). Electrophoretic analysis (Ganjeizadeh *et al*, 1995; Donnet *et al*, 2001) was also consistent with a complex of two α subunits. NKA oligomerization has been invoked to explain other experimental anomalies such as NKA ouabain-digoxin antagonism (Boldyrev, 2001; Clarke & Kane, 2007; Song *et al*, 2014a). That is, similar pharmacological inhibitors may show opposing effects that could be explained by functional interactions of protomers of a multimeric transporter complex. Functional dimerization has also been observed for other members of P-type ATPase family (Chan *et al*, 2010; Okkeri *et al*, 2002; Sackett & Kosk-Kosicka, 1996; Kosk-Kosicka *et al*, 1989; Kosk-Kosicka & Bzdega, 1988; Kanczewska *et al*, 2005; Levi *et al*, 2002; Gabizon & Friedler, 2014), including the closely related ion pump, sarco-endoplasmic reticulum Ca^2+^-ATPase (SERCA) (Bovo *et al*, 2020; Blackwell *et al*, 2016; Andersen, 1989). Kinetic studies of that transporter revealed that conformational coupling within the dimer complex enhances overall enzyme turnover by linking an energetically unfavorable step of one SERCA protomer to an energetically favorable step of the other protomer (Bovo *et al*, 2020; Froehlich *et al*, 1997). Whether this biochemical mechanism is common to other oligomeric transporters is unknown.

In this study, we probed the stoichiometry of human NKA-PLM regulatory complex using fluorescence spectroscopy and computational simulations to evaluate alternative arrangements of subunits. The results provide insight into the quaternary architecture of the complex, with implications for a structural mechanism of conformational coupling of the catalytic subunits.

## Results

### The quaternary structure of NKA-PLM regulatory complex

To investigate the architecture of the NKA αβ-FXYD complex, we fused mCyRFP1 (Bovo *et al*, 2020; Laviv *et al*, 2016; Schaaf *et al*, 2018) (FRET donor) to the N-terminus of NKA α subunit, fused mMaroon1 (Schaaf *et al*, 2018) (FRET acceptor) to the C-terminus of PLM (on the cytoplasmic side), and expressed the fusion constructs with unlabeled β1 subunit in HEK293-AAV cells. We obtained a heterogenous population of cells with varying donor/acceptor ratios and a range of fluorescence intensities. FRET was detected with fluorescence lifetime imaging microscopy (FLIM). **Fig.1A** shows cells expressing both the donor and acceptor (D+A) were characterized by a significantly shortened fluorescence lifetime (τ ~2 ns) compared to cells expressing only the donor fluorophore (D). The latter had a lifetime of 3.5 ns, similar to cells transfected with mCyRFP1-α_1_ alone (**Sup. Fig. 1A**). The data are consistent with shortening of τ by FRET. For a more detailed analysis of the fluorescence decay, we performed TCSPC point measurements, parking the excitation beam at a single spot on the plasma membrane to collect ~10^6^ photons for each record, as previously described (Blackwell *et al*, 2016; Bovo *et al*, 2020). Cells expressing mCyRFP1 (non-fusion) showed a single exponential decay with a characteristic decay time (τ) of 3.51 ns (**Sup. Fig. 1B, black**), and mCyRFP1-α_1_ was also well-described by a single exponential decay (**Fig. 1B**, black trace, “D”) with a τ of 3.52 ns (**Fig. 1C**, black points, “D”) (see Methods). Co-expression of PLM-mMaroon with mCyRFP1-α_1_ decreased the mCyRFP1-α_1_ lifetime, yielding a more complex decay (**Fig. 1B**, red, “D+A”) with average lifetime (τ_avg_) values ranging from 1-3 ns (**Fig. 1C**, red, “D+A”). α-PLM FRET was not affected by 10 μM ouabain (**Fig. 1C**, green, “D+A+OUA”). To determine whether α-PLM FRET was due to a specific interaction of donor-labeled α_1_ with acceptor-labeled α_1_, we co-expressed an excess of unlabeled α_1_. This resulted in a significant increase in τavg (**Fig. 1C**, blue “D+A+comp”). This supports the conclusion that the observed mCyRFP1-mMaroon FRET is due to a specific physical interaction of the α_1_ and PLM to which the fluorescent tags are fused. We observed similar results for α_2_ and α_3_ (**Sup. Fig. 2A-C**).

**Fig. 1.**
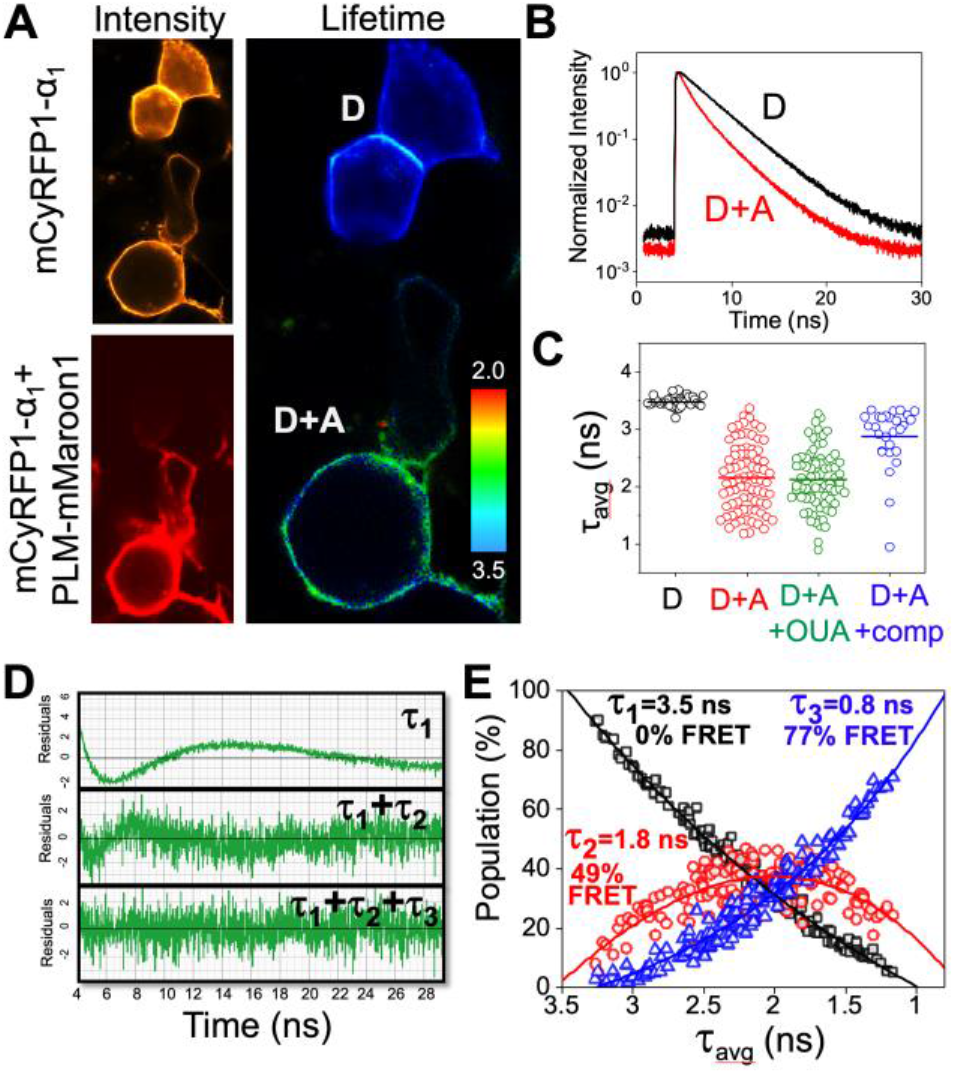
Quantification of the α_1_-PLM interaction using TCSPC. **(A)** Fluorescence intensity and fluorescence lifetime data acquired by FLIM. Cells expressing only mCyRFP1-α_1_ (donor, D) were characterized by a long lifetime. Cells coexpressing PLM-mMaroon1 (donor+acceptor, D+A) showed shorter lifetimes. **(B)** Fluorescence decay curves for donor alone (D) and donor+acceptor (D+A). **(C)** Comparison of amplitude-weighted average fluorescence lifetimes obtained by multi-exponential fitting of decay data. The donor lifetime (black, D) was shortened by FRET with the acceptor (red, D+A). The addition of 10 μM ouabain (green, OUA) did not affect detected FRET. FRET was decreased by competition with unlabeled PLM (blue, D+A+comp). **(D)** A plot of residuals of exponential decay fitting showed the fit was improved by addition of second and third components, suggesting 3 fluorescent species with distinct lifetimes. **(E)** The relative population of the three decay components depended on the overall average lifetime of the decay. Those components are attributed 3 species with FRET efficiencies of 0% (black), 49% (red), and 77% (blue).

**Fig. 2.**
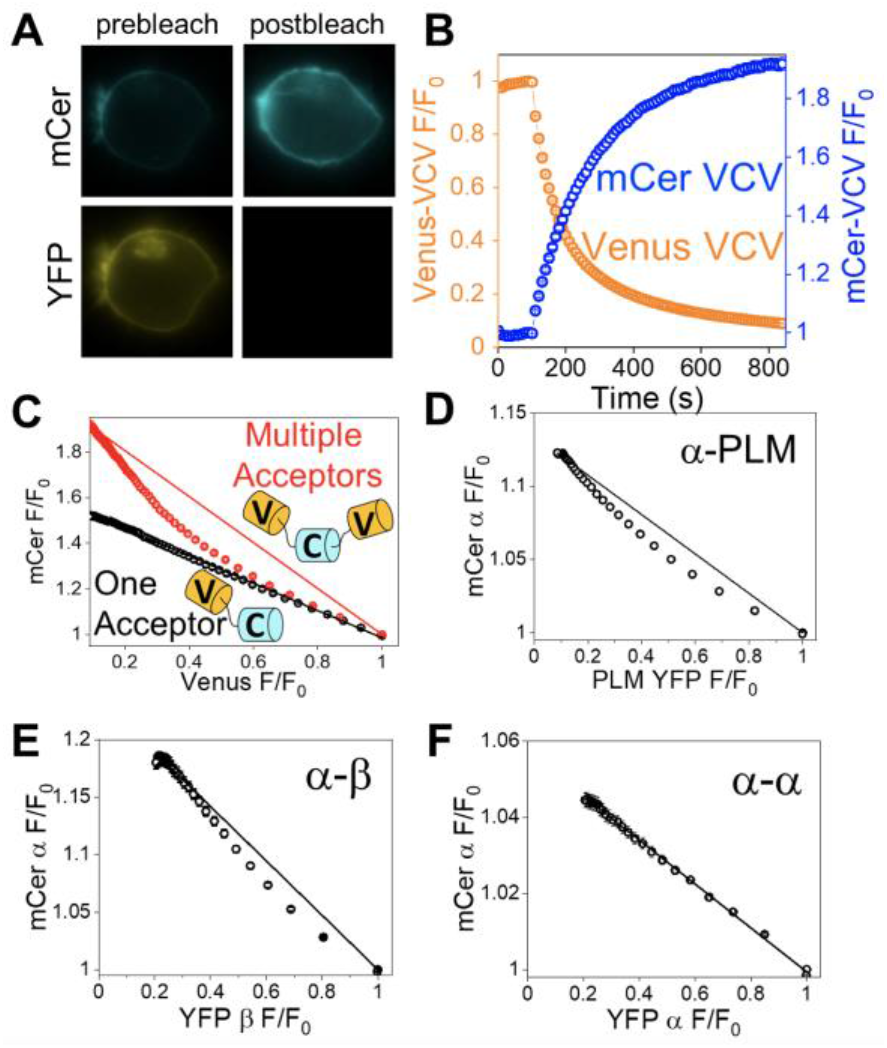
Progressive acceptor photobleaching provides insight into regulatory complex stoichiometry. **(A)** Images of cells expressing NKA and PLM labeled with mCerulean (mCer) and YFP, respectively, before and after acceptor selective photobleaching. **(B)** In a control experiment, donor (mCer) fluorescence intensity increased as acceptor (Venus) was progressively photobleached. **(C)** Replotting data (as in **B**) reveals the relationship between donor and acceptor fluorescence intensity. The relationship is linear when one acceptor is present, e. g. mCer separated from Venus by 5 amino acids (C5V, black) but is curved when there are multiple acceptors participating in FRET (VCV, red). **(D)** The regulatory complex contains multiple acceptor-labeled PLM in complex with donor-labeled α. **(E)** The complex contains multiple acceptor-labeled β. **(F)** The α-α complex produced a linear donor/acceptor relationship, suggesting a single acceptor-labeled α in complex with donor-labeled α.

The more complex decay of the D+A expressing cells was not well described by a single exponential function, as is evident from a plot of the fit residuals (**Fig. 1D**, τ_1_). Addition of a second and a third component improved the quality of the fit (**Fig. 1D**), but adding a 4^th^ decay component did not appreciably improve the residuals plot or reduced chi-squared value. Thus, the data support the existence of three species with well-resolved fluorescence lifetimes: τ_1_ = 3.46 ns, which we attribute to donors not interacting with acceptors; τ_2_ = 1.8 ns, which suggests a population of donors undergoing FRET with 49% efficiency; τ_3_ = 0.8 ns, which corresponds to a high FRET population (77 % FRET). The relative populations of these species varied from cell to cell, and we noted increased contribution of high FRET species in brightly fluorescent cells expressing a high concentration of protein. **Figure 1E** shows the results of exponential fitting of decays obtained from a survey of ~200 cells expressing a wide range of protein concentrations. The amplitude of each component lifetime (τ_1_, τ_2_, and τ_3_) varied systematically with the value of the overall decay lifetime (τ_avg_). Cells yielding decays with longer average lifetimes (τ_avg_) indicating low FRET, had strong contributions from the donor-only component (τ_1_), and low populations of the FRET species (τ_2_, τ_3_). Data from these cells falls on the left side of **Fig. 1E**. Conversely, cells manifesting short decays (low τ_avg_) had strong contributions from the high FRET species (**Fig. 1E**, blue triangles) and little contribution from the donor-alone species (black squares). Interestingly, the medium FRET species showed a biphasic dependence on τ_avg_, with low contribution to very long and very short decays, and a maximal population in decays with τ_avg_ near 2 ns. We observed the same FRET-dependent populations of species for α_1_, α_2_, and α_3_ (**Sup. Fig. 2D-F**), so data from all α isoforms are pooled and presented together in **Fig. 1E**.

To understand the nature of the 3 distinct FRET species, we sought to systematically investigate the relative population of each component as a function of protein expression. We considered that the high FRET species may represent an alternative conformation in which the donor and acceptor were in closer proximity, however, the concentration dependent appearance of this component suggests that it represents a higher-order complex of NKA and PLM containing more than one acceptor-labeled PLM. To evaluate this possibility, we performed progressive acceptor photobleaching of cells expressing mCer-NKA and PLM-YFP. This approach is illustrated in **Fig. 2A**: destruction of the YFP (acceptor) abolishes FRET, dequenching the mCer (donor). Stepwise photobleaching over time while monitoring the donor and acceptor brightness reveals the time-dependent changes in these signals (**Fig. 2B**), and plotting these signals against each other reveals the relationship between donor and acceptor fluorescence intensity (**Fig. 2C**). This donor/acceptor relationship indicates whether there is a single acceptor in the FRET complex or multiple acceptors. The standard FRET fusion construct (Kelly *et al*, 2008) C5V shows a linear plot of donor vs. acceptor brightness (**Fig. 2C**, black), while a control fusion construct with two acceptors (VCV) shows pronounced curvature (**Fig. 2C**, red) (Hou *et al*, 2008, 2012; Himes *et al*, 2016; Blackwell *et al*, 2016; Singh *et al*, 2019). This curvature occurs because FRET persists after bleaching the first acceptor in the complex, thus dequenching of the donor is delayed until later in the progressive photobleaching time course. Subjecting the mCer-NKA+PLM-YFP expressing cells to progressive acceptor photobleaching yielded a donor/acceptor relationsip with significant curvature (**Fig. 2D**), indicating that the donor-labeled NKA is in a complex with more than one acceptor-labeled PLM. The discussion section evaluates this noteworthy observation in the context of previous similar studies.

To develop a fuller picture of the NKA regulatory complex subunit stoichiometry, we engineered an acceptor-labeled β subunit and co-expressed it with donor-labeled α subunit. Progressive acceptor photobleaching yielded a non-linear D/A plot (**Fig. 2E**), suggesting the NKA complex contains multiple β subunits. Finally, we tested whether the complex included multiple α subunits by quantifying FRET from mCer-α_1_ to YFP-α_1_. We performed progressive acceptor photobleaching, and observed a linear relationship (**Fig. 2F**) suggestive of a single acceptor-labeled α protomer in complex with the donor-labeled α protomer. Collectively, these data suggest that the NKA complex includes an α-α homodimer decorated with multiple β subunits and multiple FXYD proteins. The discussion section describes how this insight into α:β:FXYD stoichiometry may be used to discriminate between alternative models of the regulatory complex quaternary architecture. In contrast to other reports that indicated de-oligomerization of NKA in the presence of the NKA inhibitor ouabain(Song *et al*, 2014a), we did not see a decrease in FRET in the presence of 10 µM ouabain, whether FRET was measured by acceptor sensitization FRET or TCSPC (**Sup. Fig. 2A-C**). Co-expression of unlabeled PLM did not increase or decrease α-α FRET measured by TCSPC (**Sup. Fig. 3B**).

**Fig. 3.**
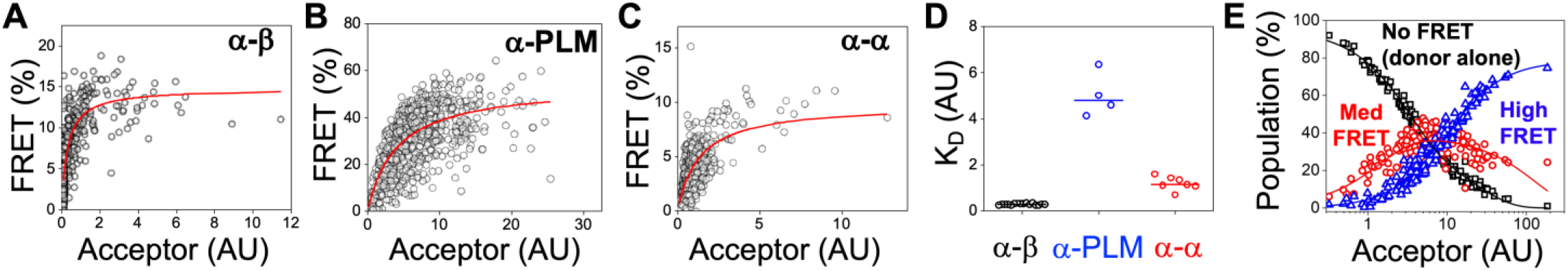
Quantification of binding affinity with acceptor sensitization FRET. **(A)** The α-β FRET binding curve revealed a very high affinity interaction. **(B)** The apparent affinity of α for PLM was lower. **(C)** α-α binding affinity was intermediate. **(D)** Summary of measured dissociation constants for α-β (black), α-PLM (blue) and α-α (red) interactions. **(E)** Indexing lifetime data from Fig. 1E with acceptor sensitization FRET measurements in Fig. 3B reveals concentration-dependent assembly of distinct FRET species.

The apparent concentration-dependent appearance of different FRET species motivated a comparison of the relative affinities of the interaction of the NKA α subunit with other components of the complex. We quantified FRET from a heterogeneous population of cells using the acceptor sensitization method, and also quantified acceptor fluorescence emission as an index of protein concentration in the cell, as previously described (Abrol *et al*, 2015, 2014; Hou & Robia, 2010; Bossuyt *et al*, 2009; Himes *et al*, 2016; Bovo *et al*, 2020; Singh *et al*, 2019). A plot of observed FRET vs. the cell acceptor fluorescence emission revealed the concentration dependence of FRET, which was well described by a hyperbolic function of the form FRET_observed_ = (FRET_max_)[protein]/K_D_+[protein], where FRET_max_ is the maximal FRET obtained at the highest concentrations and K_D_ is the apparent dissociation constant, the protein concentration that yielded half-maximal FRET (Singh *et al*, 2019). The concentration-dependence of FRET from donor labeled α subunit to acceptor labeled β suggested a very avid interaction (**Fig. 3A**) with an apparent dissociation constant that was too low to quantify reliably (K_D_ < 0.5 arbitrary units (AU)). We conclude that the α-β interaction is constitutive under these experimental conditions. The FRET based binding curve for α and PLM revealed a lower affinity interaction (**Fig. 3B**), with a higher apparent K_D_ (5.0 ± 0.8 AU). Importantly, the PLM-binding affinity is underestimated in this assay because PLM forms tetramers that do not interact with the α subunit (Song *et al*, 2011). Thus, the true concentration of monomeric PLM that is available to bind α is somewhat less than what would be estimated from total YFP-PLM fluorescence intensity. We also observed FRET from donor-labeled α to acceptor labeled α (**Fig. 3C**), with an apparent K_D_ of 1.2 ± 0.3 AU, consistent with the acceptor photobleaching experiments that suggested the NKA complex contains an α-α dimer (**Fig. 2F**). The relative dissociation constants of the α subunit interactions with other subunits of the NKA complex are summarized in **Fig. 3D**.

The observed α-PLM acceptor sensitization FRET binding curve (**Fig. 3B**) revealed the relationship between protein expression and FRET efficiency. This relationship was used to index τ_avg_ values (**Fig. 1E**, x-axis values) to protein concentration after converting τ_avg_ to observed FRET according to the relationship FRET_observed_ = 1-τ_avg_/τ_D_. Replotting of the data of **Figure 1E** as a function of relative protein concentration revealed that the NKA-PLM interaction is low in the lowest expressing cells, increasing as the protein concentration grows (**Fig. 3E**). The non-FRET species (**Fig. 3E**, black squares, τ_1_) that predominates at low protein expression levels consists of donor labeled α that is not bound to acceptor-labeled PLM. However, even at these low protein concentrations, the α-β interaction occurs, since this interaction has the highest relative affinity (**Fig. 3A**) and may be considered constitutive. At higher concentrations, the higher order complexes begin to form. In particular, the highest FRET species (**Fig. 3E**, blue triangles, τ_3_ = 0.8 ns, FRET = 77 %) is the dominant complex at the highest concentrations. Curvature analysis (**Fig. 2**) suggests this species includes multiple β subunits and multiple PLM in the complex, and it occurs at protein concentrations where α-α interactions are observed (**Fig. 3C**). The possible stoichiometry of subunits and concentration-dependent assembly pathway for the regulatory complex are discussed below.

### Hetero-dimerization of different α subunit isoforms

In this study, we primarily studied the housekeeping isoform α_1_ in evaluating NKA-PLM complex stoichiometry. To determine whether all isoforms support α-α interactions or if the interaction is specific to the α_1_ isoform, we fused mCyRFP1 and mMaroon1 to the 3 main human α isoforms and acquired FLIM data from different combinations of these α subunits. We observed the expected plasma membrane localization for the labeled proteins (**Fig. 4A-G**), and we noted a decreased donor fluorescence lifetime for every combination of isoforms (**Fig. 4H**), consistent with FRET between donor- and acceptor-labeled α subunits (**Sup. Fig. 1B, red**). Thus, α-α interactions appear to be a general feature of all three α isoforms.

**Fig. 4.**
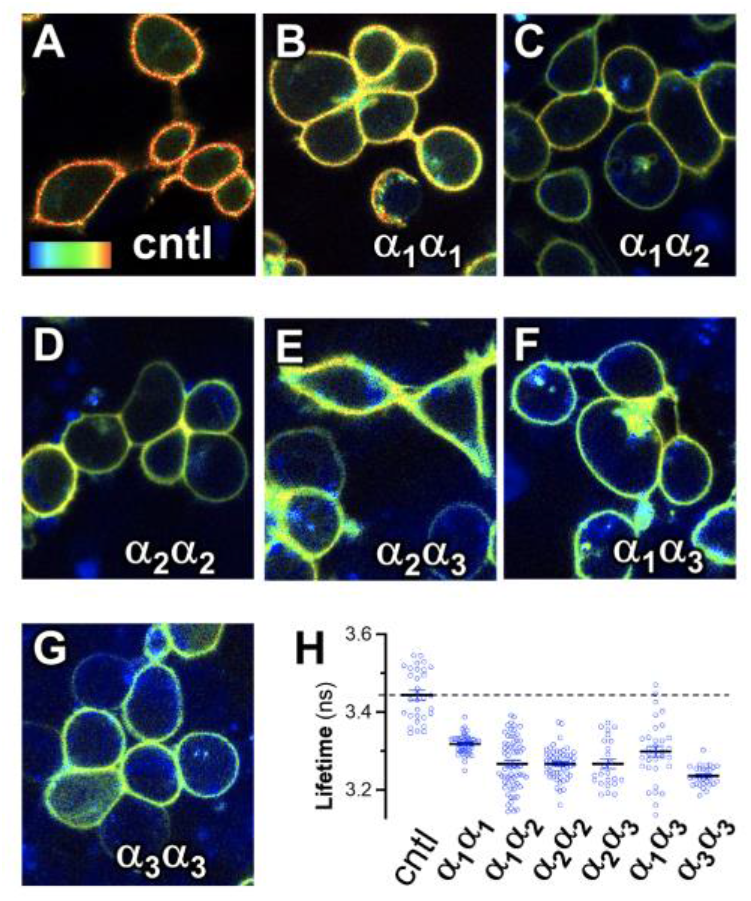
FLIM analysis of α-α FRET. **(A)** Control (donor alone). **(B-G)** Combinations of α subunit isoforms. **(H)** Quantification of FLIM images revealed all combinations of isoforms support FRET, measured as decreased lifetime compared to donor alone control (dotted line). The data suggest α-α interactions are a general mechanism. Color scale 2.2 ns (blue) to 3.4 ns (red).

### Molecular Docking of NKA α Subunit

We performed protein-protein docking simulations to generate atomistic structures of the quaternary complex formed by NKA α-β-FXYD2. Docking simulations were performed using crystal structures of NKA that captured the E1 state of the pump (PDB: 3WGU (Kanai *et al*, 2013)). The complexes were obtained with Rosetta MPDock (Alford *et al*, 2020, 2015), generating 1000 configurations that were ranked based on the Rosetta interaction energy units. Inspection of the top ranked complexes yielded three configurations that satisfy both the correct orientation and a substantial surface of intramolecular interaction. These complexes were were minimized and subjected to molecular dynamics simulations for between 0.8 μs and 1 μs.

Analysis of the MD trajectories showed that the three dimers remained structurally stable and do not dissociate on this time scale (**Sup. Fig. 3**). We compared the distance between the N-termini of the docked α subunits with that estimated from FRET experiments to determine which model best represents the structure of the complex. We found that Structural Model 1 and 2 yield α-α distance distributions (**Fig. 5A, blue, red)** that were comparable to the distance estimated from quantification of FRET, 65 Å (**Fig. 5A, dotted line)**. The distance distribution for Structural Model 1 reflects a stable, relatively compact architecture, as illustrated by a single peak centered at 60 Å (**Fig. 5A**). Structural Model 2 sampled a range of distances that distributed between two peaks at 60 Å and 70 Å. Compared to these models, Structural Model 3 exhibited substantially longer distances between α subunit N-termini (*R*=105 Å, **Fig. 5A, green**). This distance distribution is not well-matched to apparent fluorescent probe separation distance estimated by FRET. However, the fluorescent protein tags are large and connected by a flexible linker to the α subunit N-terminus, so we cannot rule out Structural Model 3. A comparison of alternative models complex 1, complex 2, and complex 3 is provided in **Sup. Fig. 4A-F**.

**Fig. 5.**
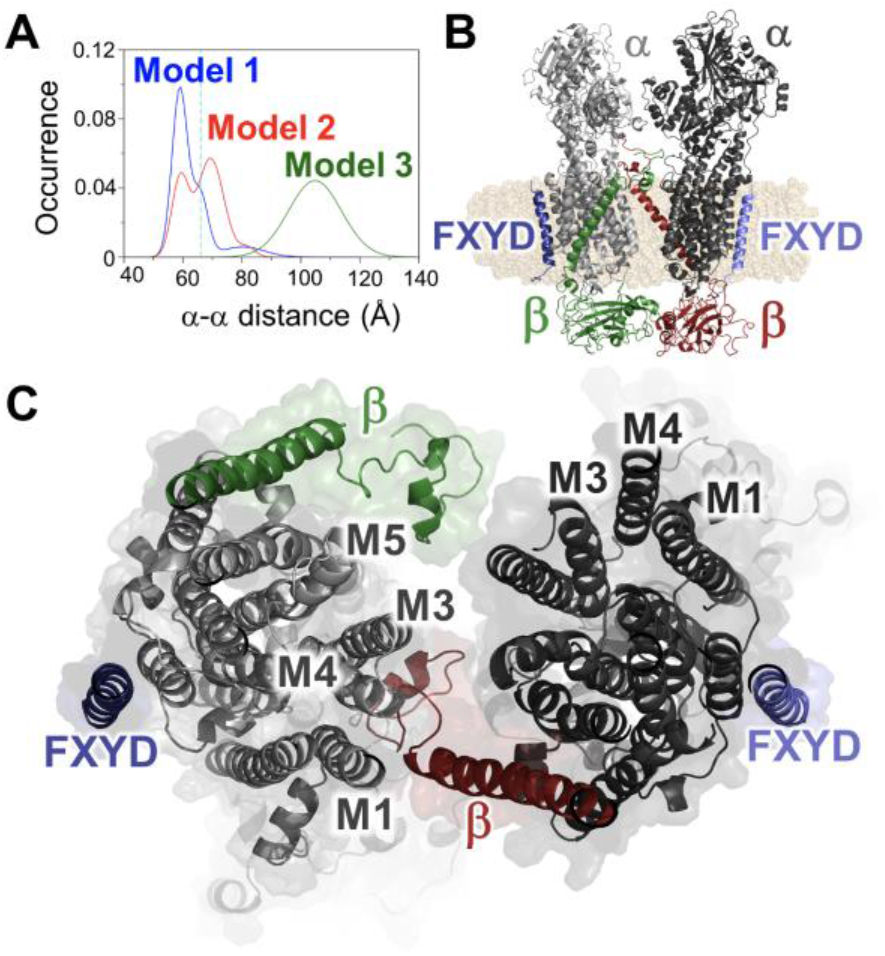
Comparing possible α-α dimer arrangements. **(A)** The distribution of distances between α subunit N-termini sampled during MD simulations. **(B)** Structural Model 1, showing docked E1P conformations of the α subunits. α subunits are shown in light gray and dark gray, β subunits are green and red; and the FXYD proteins are shown in blue and lavender. **(C)** Intramolecular contacts in Structural Model 1, viewed from the extracellular side.

The arrangement of subunits in Structural Model 1 is shown in **Fig. 5B**, with α subunit protomers shown in light and dark gray, the β subunits shown in green and red, and the FXYD proteins shown in blue and lavender. The complex is oriented with the α subunit cytoplasmic domains at the top of the figure. We noted intermolecular contacts between extracellular domains of both β subunits, and between the A- and P-domains of the α subunits. We also observed contacts between the N-terminus of the β subunit and and transmembrane helix 3 (M3) of each adjacent α subunits. These contacts examined in more detail in **Fig. 5C**, which shows a section through the center of the complex viewed from the extracellular side.

## Discussion

### The architecture of the NKA regulatory complex

Previous studies employing diverse experimental approaches have provided evidence of α subunit oligomerization in the NKA regulatory complex (Boldyrev, 2001; Blackwell *et al*, 2016; Song *et al*, 2014a; Singh, R. B., and Dhalla, 2010; Moller *et al*, 1980; Vanderkooi *et al*, 1977; Clarke & Kane, 2007). Here, we explored the binding affinity between NKA subunits and investigated the stoichiometry of the NKA regulatory complex. We considered several possible alternative stoichiometries for the regulatory complex, depicted in the schematic diagram of **Figure 6A-E**. Possible arrangements include: Stoichiometry Model **A**, a simple complex containing a single protomer of each type (αβ-PLM); **B**, αβ binds two PLM proteins at two distinct sites (high and low affinity) (PLM-αβ-PLM); **C**, αβ binds a PLM tetramer (Himes *et al*, 2016; Song *et al*, 2011; Wong *et al*, 2008; Beevers & Kukol, 2006) (αβ-PLM_4_); **D**, two αβ complexes bind to each other, along with regulatory PLM partners (αβ-PLM)_2_; and **E**, which is higher order oligomerization/aggregation (Song *et al*, 2014b). The TCSPC measurements (**Fig. 1E**) and progressive acceptor photobleaching experiments (**Fig. 2D**) rule out the simplest model, Model A (**Fig. 6A** “αβ-PLM”). Next, progressive photobleaching quantification of α-β FRET revealed a curved D/A plot that indicates multiple β subunits in the complex (**Fig. 2E**). This observation rules out Stoichiometry Models A, B, and C, which do not have multiple β subunits. Finally, we measured α-α FRET (**Fig. 2F**) with progressive photobleaching, and observed a linear plot, which shows the regulatory complex contains two α subunits. This observation therefore rules out Stoichiometry Models A, B, C, and E. We propose that Stoichiometry Model D, characterized by the complex of two α catalytic subunits plus regulatory partners (αβ-PLM)_2_, best describes the architecture of the NKA regulatory complex.

**Fig. 6.**
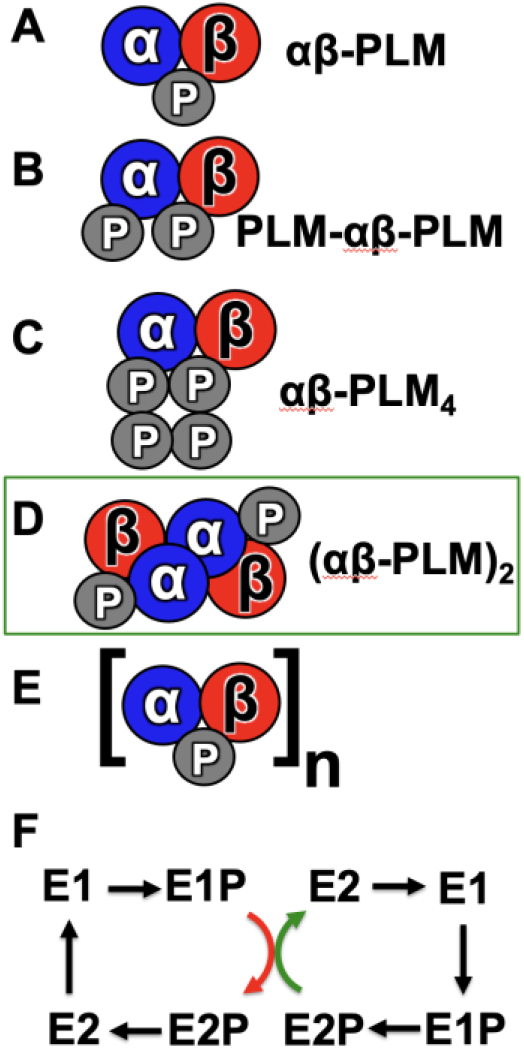
The stoichiometry, assembly, and function of the NKA regulatory complex. **(A-E)** Alternative models of NKA regulatory complex stoichiometry. The present data are consistent with Stoichiometry Model D. **(F)** A model of conformational coupling in a dimer of α subunits. An energetically favorable catalytic step of one protomer (green arrow) assists a rate limiting step (red arrow) of the other partner, enhancing overall turnover.

Complementary atomistic simulations produced several structural models of the dimer that agree with Stoichiometry Model D. The structural models all showed a high degree of symmetry (**Sup. Fig. 4**), which is a common characteristic of stable, multivalent, allosteric complexes (Goodsell & Olson, 2000). The simulations suggest an interaction between the N-termini of β and helix M3 of the α subunit, which is noteworthy because helix M3 is highly conserved across species. Mutations in this region induce changes in the functional E1-E2 transitions, alter Na^+^ binding to the transport sites, and affect the allosteric interactions that control NKA dephosphorylation (Toustrup-Jensen & Vilsen, 2002). The functional significance of this region is also emphasized by the association between mutations in the helix M3 and neurological diseases such as familial hemiplegic migraine and renal hypomagnesemia (Spadaro *et al*, 2004; Santoro *et al*, 2011; Schlingmann *et al*, 2018). We speculate that changes in α subunit functional dimerization may contribute to the deleterious effect of pathological M3 mutations.

### The Functional Significance of NKA Regulatory Complex Stoichiometry

The possibility of a dimeric NKA was first raised in 1973 (Repke & Schon, 1973; Stein *et al*, 1973), but the physiological function of the α-α interaction is still unclear. Apparent inter-protomer allosteric communication between ATP binding sites suggests a functional interaction between α subunits. Functional coupling may support synchronized cycling of pumps (Clarke *et al*, 2007; Clarke & Kane, 2007) for enhanced transport kinetics. Alternatively, the mechanism of functional coupling may follow the pattern observed for another dimeric transporter, the SERCA Ca^2+^ pump. We (Bovo *et al*, 2020; Blackwell *et al*, 2016) and others (Andersen, 1989) have observed SERCA-SERCA interactions; kinetic studies suggested that SERCA dimerization enhances Ca^2+^ transport through anti-synchronous cycling of the protomers (Mahaney *et al*, 2005). In this “conformational coupling” model, dimerization serves to speed up rate-limiting steps in the catalytic cycle of one protomer by coupling them to thermodynamically favorable steps in the other protomer (**Fig. 6F**). We hypothesize that NKA, like SERCA, may benefit from an increased catalytic efficiency and faster turnover rate due to conformational coupling of α subunit protomers.

### Concentration-Dependent Assembly of the NKA Regulatory Complex

The present progressive photobleaching experiments suggested that the regulatory complex contains more than one PLM, as shown in the pronounced curvature in the donor vs acceptor plot (**Fig. 2D**). This result was unexpected, and contrasted with our previous progressive acceptor photobleaching measurements of canine or rat NKA α_1_ in complex with mouse PLM, in which we observed only modest curvature (Bossuyt *et al*, 2009, 2006). This apparent discrepancy may be due to a lower level of protein expression that was achieved in that previous study using a CFP-NKA stable cell line. The concentration-dependent assembly of lower and higher FRET NKA complexes (**Fig. 3E**) suggests that at lower protein expression, NKA forms a simpler complex containing a single acceptor, but at higher concentrations, a larger complex containing multiple acceptor-labeled PLM subunits prevails. This larger complex with multiple acceptors gives rise to the curved D/A relationship seen in the present study **(Fig. 2D)** and accounts for the observed high FRET species seen in TCSPC experiments (**Fig. 3E**). Similarly, we have previously observed stepwise assembly of oligomers of membrane micropeptides (Singh *et al*, 2019). In that study, we found dimers formed first, yielding a linear D/A plot at low protein concentration. Then, as expression increased, the D/A curvature increased suggesting higher order oligomerization.

In considering the biological significance of the observed concentration-dependent assembly of subunits, we note possible relevance for the pathophysiology of heart failure. In transfected HEK cells, we see a range of NKA and PLM expression levels. Cells in the middle of the distribution express protein at a level that supports ~20% of the NKA population in the high FRET species (**Fig. 3E**). Comparing microsomal membrane preparations from HEK cells with membranes of cardiac myocytes suggests that NKA expression in the heart is more than 2-fold higher (**Sup. Fig. 5**). Consequently, we estimate that >60% of NKA in the heart is fully assembled in the high order complex with stoichiometry (αβ-PLM)_2_. However, under pathological conditions, such as in heart failure, NKA α subunit expression and PLM expression are significantly decreased (Bossuyt *et al*, 2005). Indeed, we observed that membrane microsomes isolated from failing human myocardium expresses NKA α at low levels that are similar to the transfected HEK cell model system used here (**Sup. Fig. 5**). Under these pathological conditions, the low protein concentration may not be sufficient to support assembly of functional regulatory complex with correct stoichiometry, (αβ-PLM)_2_.

### Summary

Collectively, our results are consistent with a sodium/potassium transporter regulatory complex that is composed of two α subunits associated with two β subunits and two PLM regulatory subunits. Docking and molecular dynamics simulations illustrated several possible quaternary arrangements of these subunits, but confirmation of the true architecture of the (αβ-PLM)_2_ regulatory complex will require high resolution structure determination under conditions that preserve the higher-order oligomeric form of NKA. We propose that α-α interactions support conformational coupling of the catalytic subunits, enhancing NKA turnover rate and transport efficiency.

## Experimental Procedures (Material and Methods)

### Molecular Biology and Cell Culture

Human NKA α_1_, β_1_ and PLM were kindly provided by Prof. Steven J. D. Karlish (Weizmann Institute of Science, Israel). α_1_ subunit encoding cDNA was subcloned into the mCerulean-C1 (mCer) modified plasmid (Addgene, USA) yielding α_1_ fused to the C-terminus of the mCer, whereas PLM was subcloned into N1-YFP vector yielding PLM fused to N-terminus of YFP. The mCyRFP1 (Laviv *et al*, 2016) vector and pcDNA3.1-mMaroon1 (Schaaf *et al*, 2018) were purchased from Addgene (Addgene, USA). Human α_1_ NKA was fused to C-terminus of mCyRFP1-C1 vector and mMaroon1 was amplified using polymerase chain reaction and inserted into N1-YFP to replace the YFP fluorescent tag. To improve NKA expression and localization, we inserted polymerase chain reaction amplificated human β_1_ subunit into unlabeled-C1 vector. All sequences have been verified by single pass primer extension analysis (ACGT Inc., USA).

HEK293-AAV cells were incubated in the humidified 5% CO2 incubator in Dulbecco’s modified Eagle’s medium (DMEM) supplemented with 10% fetal bovine serum (FBS) until 60-80% confluence. Cells were transiently transfected using Lipofectamine 3000 transfection kit (Invitrogen, USA) and transferred into DMEM supplemented with 10% FBS and 2% dimethyl sulfoxide (DMSO). 48 hours post-transfection, cells were trypsinated, plated into sterile glass bottom chamber slides and incubated at least 1 hour before imaging. The media in the chamber slides was replace by phosphate buffered saline (PBS) immediately before imaging.

### Progressive Acceptor Photobleaching

48 hours post-transfection, cells were trypsinated and plated at 200 000 cells/well density into 2-well glass bottom chamber slides and incubated for 1 hour at 37°C. Immediately before imaging, cells were washed with PBS and images of acquired by Nikon Eclipse Ti2 inverted microscope with 1.49 Apo TIRF 100x oil immersion objective with Nikon PFS system. A sequence of CFP and YFP images of the field of cells was imaged every 10 s before YFP photobleaching was applied. Acceptor was selectively photobleached using 30 s YFP exposure time and YFP, CFP sequence was recorded after each photobleaching step from total of 50 bleaching steps.

### Time Correlated Single Photon Counting (TCSPC) and Fluorescence Lifetime Imaging Microscopy (FLIM)

Time-correlated single photon counting data acquisition was performed as previously described (Blackwell *et al*, 2016; Singh *et al*, 2019). TCSPC histograms were obtained using HEK293-AAV cells expressing mCyRFP1-α_1_ NKA alone or co-expressing mCyRFP-α_1_ NKA with PLM-mMaroon1 as described in the experimental protocol. Fluorescently labeled proteins were manually selected using 500 mm focal length plan-convex lens in a flip mount to defocus the excitation supercontinuum laser beam (Fianium) to excite the whole cell. To excite mCyRFP1, the excitation laser beam was filtered through 482/18 nm bandpass filter and 0.3 neutral density filter, and emitted fluorescence was detected using 640/50 nm bandpass filter. After selection of a cell for spectroscopy, the defocusing lens was removed from the light path, the laser intensity was attenuated with a 1.0 neutral density filter, and the laser focus was positioned on a region of the cell that yielded a fluorescence intensity of 100,000 photons/s. Under these excitation conditions, we observed less than 5% photobleaching during the acquisition period. Fluorescence was detected through a 1.2 N.A. water-immersion objective with avalanche photodiode and photon counting module (PicoHarp300, PicoQuant, Germany) using a time channel width of 16 ps. 60 s of acquisition was performed for each cell, excluding cells that showed significant fluorescence intensity changes due to movement. Protein expression did not correlate well to photon count rate, which was determined primarily by how well the focused beam superimposed the plasma membrane, rather than the density of proteins in the bilayer. In several previous studies (Pallikkuth *et al*, 2013; Blackwell *et al*, 2016; Bovo *et al*, 2020; Singh *et al*, 2019) we circumvented this issue by defocusing the excitation laser to illuminate a larger portion of the microscopic field so that we could measure the fluorescence intensity of the whole cell with a camera as an index of protein expression level (Blackwell *et al*, 2016). This was less effective in the present study, as illuminating NKA required positioning of the focused beam on the plasma membrane, thus the rest of the cell was inconsistently positioned in the defocused area of illumination, compromising intensity measurements. Under these conditions, the donor-only decay was single-exponential (χ^2^ ranging from 0.961 to 1.059 for mCyRFP1-α_1_). The mCyRFP1-α_1_ construct showed a minor second component, but the amplitude of this component was only 5% so we considered it justifiable to approximate with a single exponential decay. In cells expressing donor- and acceptor-labeled proteins, additional shorter lifetimes were observed, which we attributed to FRET. Fluorescence decay histograms were analyzed using global analysis in SymPhoTime 64 software with fixed lifetime for donor alone (3.46 ns) and two freely variable global lifetimes and all variable amplitudes for all lifetimes. Analysis of distribution of acceptors in NKA-PLM complexes in **Fig. 3E** was performed using global analysis with shared lifetime and freely variable amplitude for FRET species. We observed no differences for α subunit isoforms, so the data were combined and analyzed collectively. Distances corresponding to obtained lifetimes were calculated assuming an orientation factor (κ^2^) of 2/3, and using the relationship R=R_0_(E^−1^-1)^1/6^ (Forster, 1948), where R_0_ is the Förster distance for mCyRFP1-mMaroon1 pair (63.34 Å) (Schaaf *et al*, 2018) and E is average FRET efficiency from lifetime measurements.

Relative protein concentrations were interpolated from maximal FRET efficiency determined using TCSPC with mCyRFP1 and mMaroon1 and FRET_max_ obtained from hyperbolic fit of acceptor sensitization FRET binding curve of constructs labeled with CFP and YFP. The mMaroon1 protein concentration was calculated using following relationship [mMaroon1]=Average FRET.K_D_/(FRET_max_-Average FRET), where average FRET is equal to 1-(τ_DA_/τ_D_), τ_DA_ is lifetime of donor in the presence of acceptor, τ_D_ is lifetime of donor alone, FRET_max_ is value of maximal FRET efficiency detected for all cells in the dataset (74.47% in our case) and K_D_ is dissociation constant.

### Statistics

Data are represented as mean ± standard error of the mean (SEM) of n measurements. Statistical comparison between two groups were performed in OriginPro2020b software (OriginLab) using Student’s t-test for unpaired data sets. The differences considered as statistically significant had p<0.05.

### Acceptor Sensitization Measurements

This experiment was performed as previously described in (Bidwell *et al*, 2011; Singh *et al*, 2019) with some modifications. Similar to progressive acceptor photobleaching experiment, cells were trypsinated 48 hours post-transfection and plated at 100 000 cells/well density into 4-well glass bottom chamber slides and incubated for 1 hour at 37°C. Subsequently, cells were washed twice with PBS, followed by labelling of the cell plasma membrane using 2 μg/ml wheat germ agglutinin with Alexa Fluor 594 (Molecular Probes, USA). Cells were incubated for 5 minutes at room temperature and access of WGA was washed away by three washes using PBS. Images of cells were acquired by Nikon Eclipse Ti2 inverted microscope with NA 40x air objective. Acceptor sensitized FRET was used to measure binding affinity of mCer3-NKA and PLM-eYFP constructs. For each field, the set of four images of CFP, YFP, CFP-YFP FRET and WGA Alexa Fluor 594 fluorescence were acquired using following exposure times of 500 ms, 100 ms, 500 ms and 50 ms, respectively. Specific fluorophore was excited with adequate excitation wavelength with subsequent emission detection for CFP, YFP and WGA channels with exception of CFP-YFP FRET where excitation wavelength for CFP was used and YFP emission was captured. Then stage was shifted to a new position and another field was imaged without overlapping areas. To ensure that cells are remaining in focus, Nikon PFS feedback system was used.

Each field of the WGA Alexa Fluor 594 image was used to create a mask representing only membrane localized proteins. This mask was subsequently applied to each channel for selected field of cells.

### Molecular Docking of NKA Dimers

To explore the atomic details of the NKA dimer interface, we performed molecular docking of NKA dimers in different states of the reaction cycle using the following crystal structures: 3KDP (E2P) (Morth *et al*, 2007), 7DDJ (E2Pi) (Kanai *et al*, 2020), and 3WGU (E1Pi.ADP) (Kanai *et al*, 2013). Docking was performed following the strategy described in (Alford *et al*, 2020), in search of interactions between either homo- or heterodimers (with 3KDP (Morth *et al*, 2007) in the E2P state). Specifically, we used ClusPro for global docking to generate possible starting structures. Following that, we used Rosetta MPDock for local refinement of these structures.

The ClusPro web server (Kozakov *et al*, 2017, 2013; Desta *et al*, 2020; Vajda *et al*, 2017) was first used to perform global docking of NKA dimers using default settings. The starting structures were stripped of hetero atoms and only the protein chains were used. Around 100 resulting structures were then screened based on: 1) whether the monomers are in parallel alignment (i.e., the angle between the monomers should not exceed 40°), and 2) whether the transmembrane domains of each monomer were located within the same plane (i.e., the distance between the transmembrane regions of the monomers should not exceed 5 Å). Dimer pairs that met the above criteria were next subject to local refinement using Rosetta MPDock, a docking algorithm that runs protein-protein docking in the membrane bilayer (Alford *et al*, 2015; Gray *et al*, 2003; Chaudhury *et al*, 2011). The general Rosetta MPDock protocol consists of three steps: 1) generate a membrane span file that defines the transmembrane regions of the proteins; 2) prepack the docking partners to optimize side chain orientation; and 3) protein-protein docking in the membrane environment. As a constantly updated database of transmembrane proteins from PDB, PDBTM provides not only protein coordinates but also transmembrane profiles of the proteins (Tusnády *et al*, 2004, 2005; Kozma *et al*, 2013). The span files for the crystal structures were therefore generated based on PDBTM coordinates files with the exception of 7DDJ, for which such files were not available. Accordingly, the transmembrane region of 7DDJ was defined by using the transmembrane domain from 3KDP as they share similar structures in that domain (RMSD = 1.506 Å). Detailed input span files and flags used are provided in the Supplementary materials. For each dimer pair, Rosetta MPDock generated 1,000 structures. These 1,000 structures were ranked by the Rosetta interaction energy (I_sc). We compared best 10 conformations with our experimental data and selected 3 candidates with best agreement between docked structures and experimental measurements. All those 3 candidates represented 3WGU-3WGU combination and were used as initial models for molecular dynamic simulations.

### Molecular dynamic simulations

We modeled NKA residues Glu327, Glu779, Asp804, and Glu954 as protonated (Rui *et al*, 2016) to preserve the structural stability of the E1 state. The complexes were inserted in a pre-equilibrated 150 × 150 Å bilayer of POPC:POPE:POPS lipids (2:1:1) to mimic the composition of the plasma membrane. In this study, we used the replacement method to generate lipid packing around the αβ-PLM dimers. The initial systems were solvated using TIP3P water molecules with a minimum margin of 25 Å in the z-axis between the edges of the periodic box and the protein. Na^+^, K^+^ and Cl^−^ ions were added to neutralize the system and to produce Na^+^ and K^+^ concentrations of 140 and 5 mM, respectively (Čechová *et al*, 2016; Zacchia *et al*, 2016). Molecular simulations were performed with AMBER20 on Tesla V100 GPUs (Salomon-Ferrer *et al*, 2013) using the AMBER ff19SB force field (Tian *et al*, 2020). We maintained a temperature of 310 K with a Langevin thermostat, and a pressure of 1.0 bar with the Monte Carlo barostat. We used the SHAKE algorithm to constrain all bonds involving hydrogens and allow a time step of 2 fs. The systems were subjected to 5000 steps of steepest-descent energy minimization followed by equilibration as follows: two 25-ps MD simulations using a canonical ensemble (NVT), one 25-ps MD simulation using an isothermal–isobaric ensemble (NPT), and two 5-ns MD simulations using the NPT ensemble. The equilibrated were used as a starting point to perform a single MD trajectory of each complex for a total simulation time of 2.6 μs using an NPT ensemble.

## Author Contributions

J. S. performed FLIM, TCSPC, progressive acceptor photobleaching and acceptor sensitization FRET experiments in the study, analyzed data and wrote the manuscript. N. R. H. performed and analyzed isoform specific FLIM data. J. Q. Y. performed western blotting and reviewed manuscript. S. L. C helped with analysis and reviewed the manuscript. X. F. performed molecular docking. P. M. K.-H. designed molecular docking study. L. M. E.-F. performed, analyzed, and wrote the section about the molecular dynamic simulations of dimer complexes and reviewed the manuscript. J. B. B. helped design the study, interpret the results, and review the manuscript. S. L. R. supervised the project and helped to write the manuscript.

## Acknowledgments

The authors are grateful to the patients and organ donors who donated the samples used in this project (IRB 210940821918). The authors thank Steven J. Karlish for DNA encoding human NKA and PLM; Petra Cechova, Greg Gillispie, and Pablo Artigas for helpful discussions; Ellen E. Cho and Marsha P. Pribadi for technical assistance. This research was supported by American Heart Association Postdoctoral Fellowship 830562 (to J.S.), the Loyola Cardiovascular Research Institute, and the National Institute of Health grants HL092321 and HL143816 (to S. L. R.), R35 GM124977 (to P.M. K.-H.), R01GM120142 (to L. M. E.-F.), and R01HL142282-04 (to J. B. B.) This research was supported in part through computational resources and services provided by Advanced Research Computing at the University of Michigan, Ann Arbor, Michigan.

## REFERENCES

Abrol N, Smolin N, Armanious G, Ceholski DK, Trieber CA, Young HS & Robia SL (2014) Phospholamban C-terminal residues are critical determinants of the structure and function of the calcium ATPase regulatory complex. J Biol Chem 289: 25855–25866

Abrol N, De Tombe PP & Robia SL (2015) Acute inotropic and lusitropic effects of cardiomyopathic R9C mutation of phospholamban. J Biol Chem 290: 7130–7140

Alford RF, Koehler Leman J, Weitzner BD, Duran AM, Tilley DC, Elazar A & Gray JJ (2015) An Integrated Framework Advancing Membrane Protein Modeling and Design. PLoS Comput Biol 11: e1004398

Alford RF, Smolin N, Young HS, Gray JJ & Robia SL (2020) Protein Docking and Steered Molecular Dynamics Reveal Alternative Regulatory Sites on the SERCA Calcium Transporter. 1–18

Andersen JP (1989) Monomer-oligomer equilibrium of sarcoplasmic reticulum Ca-ATPase and the role of subunit interaction in the Ca2+ pump mechanism. Biochim Biophys Acta 988: 47–72

Askari A (1987) (Na++K+)-ATPase: on the number of the ATP sites of the functional unit. J Bioenerg Biomembr 19: 359–374

Beevers AJ & Kukol A (2006) Secondary structure, orientation, and oligomerization of phospholemman, a cardiac transmembrane protein. Protein Sci 15: 1127–1132

Bidwell P, Blackwell DJ, Hou Z, Zima A V. & Robia SL (2011) Phospholamban binds with differential affinity to calcium pump conformers. J Biol Chem 286: 35044–35050

Blackwell DJ, Zak TJ & Robia SL (2016) Cardiac Calcium ATPase Dimerization Measured by Cross-Linking and Fluorescence Energy Transfer. Biophys J 111: 1192–1202

Boldyrev AA (2001) Na/K-ATPase as an oligomeric ensemble. Biochem 66: 1013–1025

Bossuyt J, Ai X, Moorman JR, Pogwizd SM & Bers DM (2005) Expression and phosphorylation of the Na-pump regulatory subunit phospholemman in heart failure. Circ Res 97: 558–565

Bossuyt J, Despa S, Han F, Hou Z, Robia SL, Lingrel JB & Bers DM (2009) Isoform specificity of the Na/K-ATPase association and regulation by phospholemman. J Biol Chem 284: 26749–26757

Bossuyt J, Despa S, Martin JL & Bers DM (2006) Phospholemman phosphorylation alters its fluorescence resonance energy transfer with the Na/K-ATPase pump. J Biol Chem 281: 32765–32773

Bovo E, Nikolaienko R, Cleary SR, Seflova J, Kahn D, Robia SL & Zima A V. (2020) Dimerization of SERCA2a Enhances Transport Rate and Improves Energetic Efficiency in Living Cells. Biophys J 119: 1456–1465

Čechová P, Berka K & Kubala M (2016) Ion Pathways in the Na+/K+-ATPase. J Chem Inf Model 56: 2434–2444

Chan H, Babayan V, Blyumin E, Gandhi C, Hak K, Harake D, Kumar K, Lee P, Li TT, Liu HY, et al (2010) The P-Type ATPase Superfamily. J Mol Microbiol Biotechnol 19: 5–104

Chaudhury S, Berrondo M, Weitzner BD, Muthu P, Bergman H & Gray JJ (2011) Benchmarking and analysis of protein docking performance in Rosetta v3.2. PLoS One 6

Clarke RJ, Apell HJ & Kong BY (2007) Allosteric effect of ATP on Na+,K+-ATPase Conformational kinetics. Biochemistry 46: 7034–7044

Clarke RJ & Kane DJ (2007) Two Gears of Pumping by the Sodium Pump. Biophys J 93: 4187–4196

Desta IT, Porter KA, Xia B, Kozakov D & Vajda S (2020) Performance and Its Limits in Rigid Body Protein-Protein Docking. Structure 28: 1071–1081.e3

Donnet C, Arystarkhova E & Sweadner KJ (2001) Thermal Denaturation of the Na,K-ATPase Provides Evidence for α-α Oligomeric Interaction and γ Subunit Association with the C-terminal Domain. J Biol Chem 276: 7357–7365

Forster T (1948) Zwischenmolekulare energiewanderung und fluoreszenz. Ann Phys 437: 55–75

Froehlich JP, Taniguchi K, Fendler K, Mahaney JE, Thomas DD & Albers RW (1997) Complex kinetic behavior in the Na,K- and Ca-ATPases. Evidence for subunit-subunit interactions and energy conservation during catalysis. Ann N Y Acad Sci 834: 280–296

Gabizon R & Friedler A (2014) Allosteric modulation of protein oligomerization: an emerging approach to drug design. Front Chem 2

Ganjeizadeh M, Zolotarjova N, Huang W-H & Askari A (1995) Interactions of Phosphorylation and Dimerizing Domains of the alpha subunit of Na+/K+-ATPase. J Biol Chem 270: 15707–15710

Geering K (2005) Function of FXYD proteins, regulators of Na,K-ATPase. J Bioenerg Biomembr 37: 387–392

Goodsell DS & Olson AJ (2000) Structural Symetry and Protein Function. Annu Rev Biophys Biomol Struct 29: 105–153

Gray JJ, Moughon S, Wang C, Schueler-Furman O, Kuhlman B, Rohl CA & Baker D (2003) Protein-protein docking with simultaneous optimization of rigid-body displacement and side-chain conformations. J Mol Biol 331: 281–299

Himes RD, Smolin N, Kukol A, Bossuyt J, Bers DM & Robia SL (2016) L30A mutation of phospholemman mimics effects of cardiac glycosides in isolated cardiomyocytes. Biochemistry 55: 6196–6204

Hou Z, Hu Z, Blackwell DJ, Miller TD, Thomas DD & Robia SL (2012) 2-Color calcium pump reveals closure of the cytoplasmic headpiece with calcium binding. PLoS One 7: 1–10

Hou Z, Kelly EM & Robia SL (2008) Phosphomimetic mutations increase phospholamban oligomerization and alter the structure of its regulatory complex. J Biol Chem 283: 28996–29003

Hou Z & Robia SL (2010) Relative affinity of calcium pump isoforms for phospholamban quantified by fluorescence resonance energy transfer. J Mol Biol 402: 210–216

Kanai R, Cornelius F, Ogawa H, Motoyama K, Vilsen B & Toyoshima C (2020) Binding of cardiotonic steroids to Na+,K+-ATPase in the E2P state. Proc Natl Acad Sci U S A 118: 1–12

Kanai R, Ogawa H, Vilsen B, Cornelius F & Toyoshima C (2013) Crystal structure of a Na+-bound Na+,K+-ATPase preceding the E1P state. Nature 502: 201–6

Kanczewska J, Marco S, Vandermeeren C, Maudoux O, Rigaud J-L & Boutry M (2005) Activation of the plant plasma membrane H-ATPase by phosphorylation and binding of 14-3-3 proteins converts a dimer into a hexamer. Proc Natl Acad Sci U S A 102: 11675 LP – 11680

Kelly EM, Hou Z, Bossuyt J, Bers DM & Robia SL (2008) Phospholamban oligomerization, quaternary structure, and sarco(endo)plasmic reticulum calcium ATPase binding measured by fluorescence resonance energy transfer in living cells. J Biol Chem 283: 12202–12211

Kosk-Kosicka D & Bzdega T (1988) Activation of the erythrocyte Ca2+-ATPase by either self-association or interaction with calmodulin. J Biol Chem 263: 18184–18189

Kosk-Kosicka D, Bzdega T & Wawrzynow A (1989) Fluorescence energy transfer studies of purified erythrocyte Ca2+-ATPase. Ca2+-regulated activation by oligomerization. J Biol Chem 264: 19495–19499

Kozakov D, Beglov D, Bohnuud T, Mottarella SE, Xia B, Hall DR & Vajda S (2013) How good is automated protein docking? Proteins Struct Funct Bioinforma 81: 2159–2166

Kozakov D, Hall DR, Xia B, Porter KA, Padhorny D, Yueh C, Beglov D & Vajda S (2017) The ClusPro web server for protein-protein docking. Nat Protoc 12: 255–278

Kozma D, Simon I & Tusnády GE (2013) PDBTM: Protein data bank of transmembrane proteins after 8 years. Nucleic Acids Res 41: 524–529

Laviv T, Kim BB, Chu J, Lam AJ, Lin MZ & Yasuda R (2016) Simultaneous dual-color fluorescence lifetime imaging with novel red-shifted fluorescent proteins. Nat Methods 13: 989–992

Levi V, Rossi JP, Castello PR & González Flecha FL (2002) Structural significance of the plasma membrane calcium pump oligomerization. Biophys J 82: 437–446

Mahaney JE, Albers RW, Waggoner JR, Kutchai HC & Froehlich JP (2005) Intermolecular conformational coupling and free energy exchange enhance the catalytic efficiency of cardiac muscle SERCA2a following the relief of phospholamban inhibition. Biochemistry 44: 7713–7724

Moller J V., Lind KE & Andersen JP (1980) Enzyme kinetics and substrate stabilization of detergentsolubilized and membraneous (Ca2+ + Mg2+)-activated ATPase from sarcoplasmic reticulum. Effect of protein-protein interactions. J Biol Chem 255: 1912–1920

Morth JP, Pedersen BP, Toustrup-Jensen MS, Sørensen TLM, Petersen J, Andersen JP, Vilsen B & Nissen P (2007) Crystal structure of the sodium-potassium pump. Nature 450: 1043–9

Newman RA, Yang P, Pawlus AD & Block KI (2008) Cardiac glycosides as novel cancer therapeutic agents. Mol Interv 8: 36–49

Okkeri J, Bencomo E, Pietila M & Haltia T (2002) Introducing Wilson disease mutations into the zinc-transporting P-type ATPase of Escherichia coli: The mutation P634L in the ‘hinge’ motif (GDGXNDXP) perturbs the formation of the E2P state. Eur J Biochem 269: 1579–1586

Ottolenghi P & Jensen J (1983) The K+-induced apparent heterogeneity of high-affinity nucleotide-binding sites in (Na+ + K+)-ATPase can only be due to the oligomeric structure of the enzyme. Biochim Biophys Acta 727: 89–100

Pallikkuth S, Blackwell DJ, Hu Z, Hou Z, Zieman DT, Svensson B, Thomas DD & Robia SL (2013) Phosphorylated phospholamban stabilizes a compact conformation of the cardiac calcium-ATPase. Biophys J 105: 1812–1821

Palmer CJ, Scott BT & Jones LR (1991) Purification and complete sequence determination of the major plasma membrane substrate for cAMP-dependent protein kinase and protein kinase C in myocardium. J Biol Chem 266: 11126–11130

Repke k. RH & Schon R (1973) Flip-flop model of (Na,K)-ATPase function. Acta Biol Med Germ 31: K19–K30

Rui H, Artigas P & Roux B (2016) The selectivity of the Na+/K+-pump is controlled by binding site protonation and self-correcting occlusion. Elife 5: 1–24

Sackett DL & Kosk-Kosicka D (1996) The active species of plasma membrane Ca2+-ATPase are a dimer and a monomer-calmodulin complex. J Biol Chem 271: 9987–91

Salomon-Ferrer R, Götz AW, Poole D, Le Grand S & Walker RC (2013) Routine Microsecond Molecular Dynamics Simulations with AMBER on GPUs. 2. Explicit Solvent Particle Mesh Ewald. J Chem Theory Comput 9: 3878–3888

Santoro L, Manganelli F, Fortunato MR, Soldovieri MV, Ambrosino P, Iodice R, Pisciotta C, Tessa A, Santorelli F & Taglialatela M (2011) A new Italian FHM2 family: Clinical aspects and functional analysis of the disease-associated mutation. Cephalalgia 31: 808–819

Schaaf TM, Li A, Grant BD, Peterson K, Yuen S, Bawaskar P, Kleinboehl E, Li J, Thomas DD & Gillispie GD (2018) Red-shifted FRET biosensors for high-throughput fluorescence lifetime screening. Biosensors 8: 1–15

Schlingmann KP, Bandulik S, Mammen C, Tarailo-Graovac M, Holm R, Baumann M, Konig J, Lee JJY, Drogemoller B, Imminger K, et al (2018) Germline De Novo Mutations in ATP1A1 Cause Renal Hypomagnesemia, Refractory Seizures, and Intellectual Disability. Am J Hum Genet 103: 808–816

Singh, R. B., and Dhalla NS (2010) Ischemia-reperfusion-induced changes in sarcolemmal Na+/K+-ATPase are due to the activation of calpain in the heart. Can J Physiol Pharmacol 88: 388–397

Singh DR, Dalton MP, Cho EE, Pribadi MP, Zak TJ, Šeflová J, Makarewich CA, Olson EN, Robia SL, Singh DR, et al (2019) Newly Discovered Micropeptide Regulators of SERCA Form Oligomers but Bind to the Pump as Monomers. J Mol Biol 431: 4429–4443

Song H, Karashima E, Hamlyn JM & Blaustein MP (2014a) Ouabain-digoxin antagonism in rat arteries and neurones. J Physiol 592: 941–969

Song H, Karashima E, Hamlyn JM & Blaustein MP (2014b) Ouabain-digoxin antagonism in rat arteries and neurones. J Physiol 592: 941–969

Song Q, Pallikkuth S, Bossuyt J, Bers DM & Robia SL (2011) Phosphomimetic mutations enhance oligomerization of phospholemman and modulate its interaction with the Na/K-ATPase. J Biol Chem 286: 9120–9126

Spadaro M, Ursu S, Lehmann-Horn F, Liana V, Giovanni A, Paola G, Frontali M & Jurkat-Rott K (2004) A G301R Na+/K+-ATPase mutation causes familial hemiplegic migraine type 2 with cerebellar signs. Neurogenetics 5: 177–185

Stein WD, Lieb WR, Karlish SJ & Eilam Y (1973) A model for active transport of sodium and potassium ions as mediated by a tetrameric enzyme. Proc Natl Acad Sci 70: 275–278

The Digitalis Investigational Group (1997) The effect of digoxin on mortality and morbidity in patients with heart failure. N Engl J Med 336: 525–533

Tian C, Kasavajhala K, Belfon KAA, Raguette L, Huang H, Migues AN, Bickel J, Wang Y, Pincay J, Wu Q, et al (2020) Ff19SB: Amino-Acid-Specific Protein Backbone Parameters Trained against Quantum Mechanics Energy Surfaces in Solution. J Chem Theory Comput 16: 528–552

Toustrup-Jensen MS & Vilsen B (2002) Importance of Glu 282 in Transmembrane Segment M3 of the Na +, K + - ATPase for Control of Cation Interaction and Conformational Changes. J Biol Chem 277: 38607–38617

Tusnády GE, Dosztányi Z & Simon I (2004) Transmembrane proteins in the Protein Data Bank: Identification and classification. Bioinformatics 20: 2964–2972

Tusnády GE, Dosztányi Z & Simon I (2005) PDB_TM: Selection and membrane localization of transmembrane proteins in the protein data bank. Nucleic Acids Res 33: 275–278

Vajda S, Yueh C, Beglov D, Bohnuud T, Mottarella SE, Xia B, Hall DR & Kozakov D (2017) New additions to the ClusPro server motivated by CAPRI. Proteins Struct Funct Bioinforma 85: 435–444

Vanderkooi JM, Ierokomas A, Nakamura H & Martonosi A (1977) No Title. Biochemistry 16: 1262–1267

Wong A, Beevers AJ, Kukol A, Dupree R & Smith ME (2008) Solid-state 17O NMR spectroscopy of a phospholemman transmembrane domain protein: implications for the limits of detecting dilute 17O sites in biomaterials. Solid State Nucl Magn Reson 33: 72–75

Yap JQ, Seflova J, Sweazey R, Artigas P & Robia SL (2021) FXYD proteins and sodium pump regulatory mechanisms. J Gen Physiol 153: 1–18

Zacchia M, Abategiovanni ML, Stratigis S & Capasso G (2016) Potassium: From Physiology to Clinical Implications. Kidney Dis 2: 72–79

