## Supplementary Materials for "STOICHIOMETRY OF THE SODIUM PUMP-PHOSPHOLEMMAN REGULATORY COMPLEX"

**Supplementary Methods:**

### Human Heart Tissue Procurement

Human left-ventricular tissue was provided by Loyola Cardiovascular Research Institute Biorepository (IRB number 210940821918). Informed consent was obtained for collection of heart tissue. Tissue from failing human hearts with non-ischemic idiopathic dilated cardiomyopathy was collected at the time of heart explant or at the time of LVAD implantation and flash frozen in liquid nitrogen.

### Human Tissue Membrane Protein enrichment

Frozen left-ventricular tissue was placed in 5 mL of a Buffer A (100 mM KCl, 2.5 mM $K_2HPO_4$, 2.5 mM $KH_2PO_4$, 2 mM EDTA) containing protease and phosphatase inhibitors (1:100 ratio). The tissue homogenized with a mechanical homogenizer and rotated for 1hr at 4°C. Samples were centrifuged at 6,400 g for 20 minutes at 4°C. The supernatant containing soluble proteins from the previous step was centrifuged at 10,000 × g for 20 minutes to remove heavy debris. Subsequently, the supernatant was centrifuged at 48 000 x g at 4°C. Finally, the pellet containing the membrane proteins was dissolved in 100 µl of Buffer B (1 M sucrose, 50 mM KCl) and stored at -80°C. The total protein concentration was determined using Bicinchoninic Acid (BCA) Kit for Protein Determination (Pierce), and all reagents were purchased from Sigma Aldrich if not stated otherwise.

### HEK Cell Membrane Protein enrichment

48 hours post-transfection, cells were scraped in 5 ml homogenizing buffer (HB) composed of 250 mM sucrose, 10 mM Tris, 2 mM EDTA pH 7.4 with protease inhibitor and centrifuged cells for 10 min at 4000 × g. Cell pellet was re-suspended in HB with protease inhibitors and homogenized. Lysed cells were centrifuged for 20 min at 4000 × g. Supernatant was collected and centrifuged at 55,000 × g for 30 min to collect total membrane fraction.

### Western blotting

After isolating the membrane fraction, protein concentration was determined using a BCA assay (Thermo Fisher). Samples were denatured in 4x Laemmli sample buffer with beta-mercaptoethanol at 90°C for 5 min, run on a 4–15% polyacrylamide gradient gel (Bio-Rad, Hercules, CA) and transferred to polyvinylidene difluoride membrane. . Following transfer, membranes were incubated with Revert Total Protein Stain (LI-COR Biosciences) for 5 min to obtain total protein in each lane and then blocked in a 1:1 ratio of Intercept Blocking Buffer (LI-COR Biosciences) to phosphate buffered saline with Tween (PBS-T) solution for 1 h at room temperature. Afterwards, membranes were incubated with primary antibody, anti-pan NKA from Development Studies Hybridoma Bank, University of Iowa (1:1000, cat. no. AB2166869) overnight in PBS-T at 4 °C with gentle rocking. Blots were incubated with anti-mouse secondary antibody for 1 h at room temperature (1:10,000 dilution) and analyzed using the LI-COR Image Studio software.

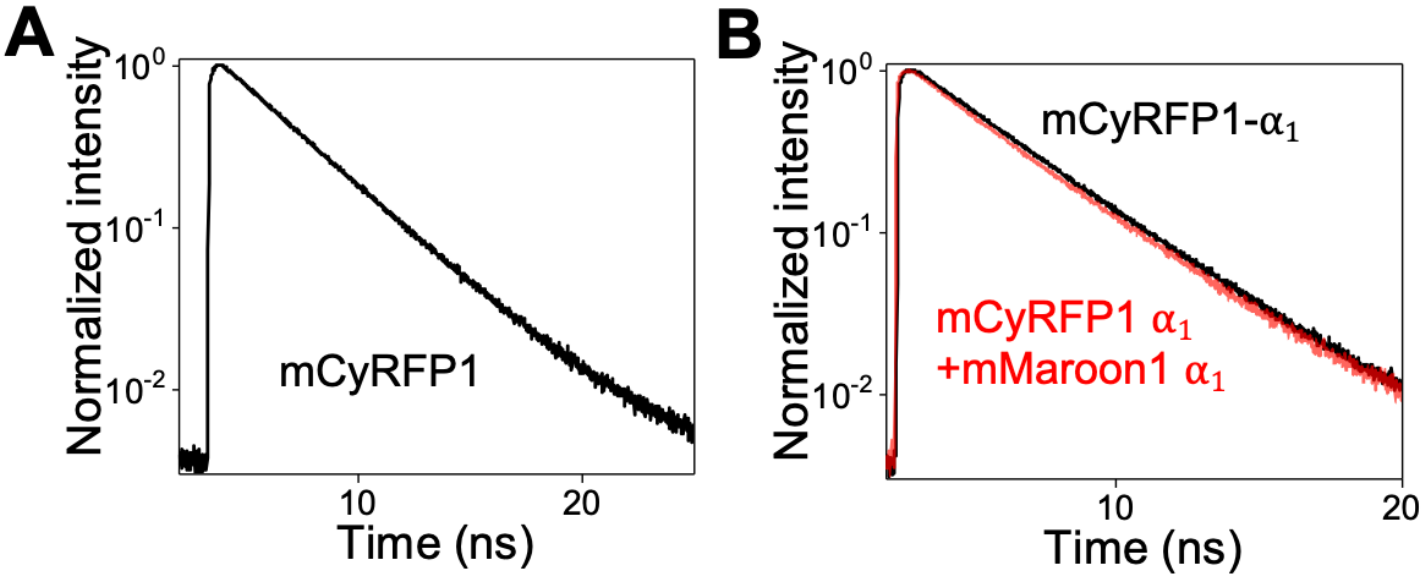


**Sup. Fig. 1**: Quantification of NKA-PLM regulatory complex using TCSPC. (A) Free (non-fusion) mCyRFP1 protein decays with single exponential kinetics. (B) The fluorescence lifetime of NKA α subunit labeled with mCyRFP1 is shortened by FRET with mMaroon1-labeled α subunit.

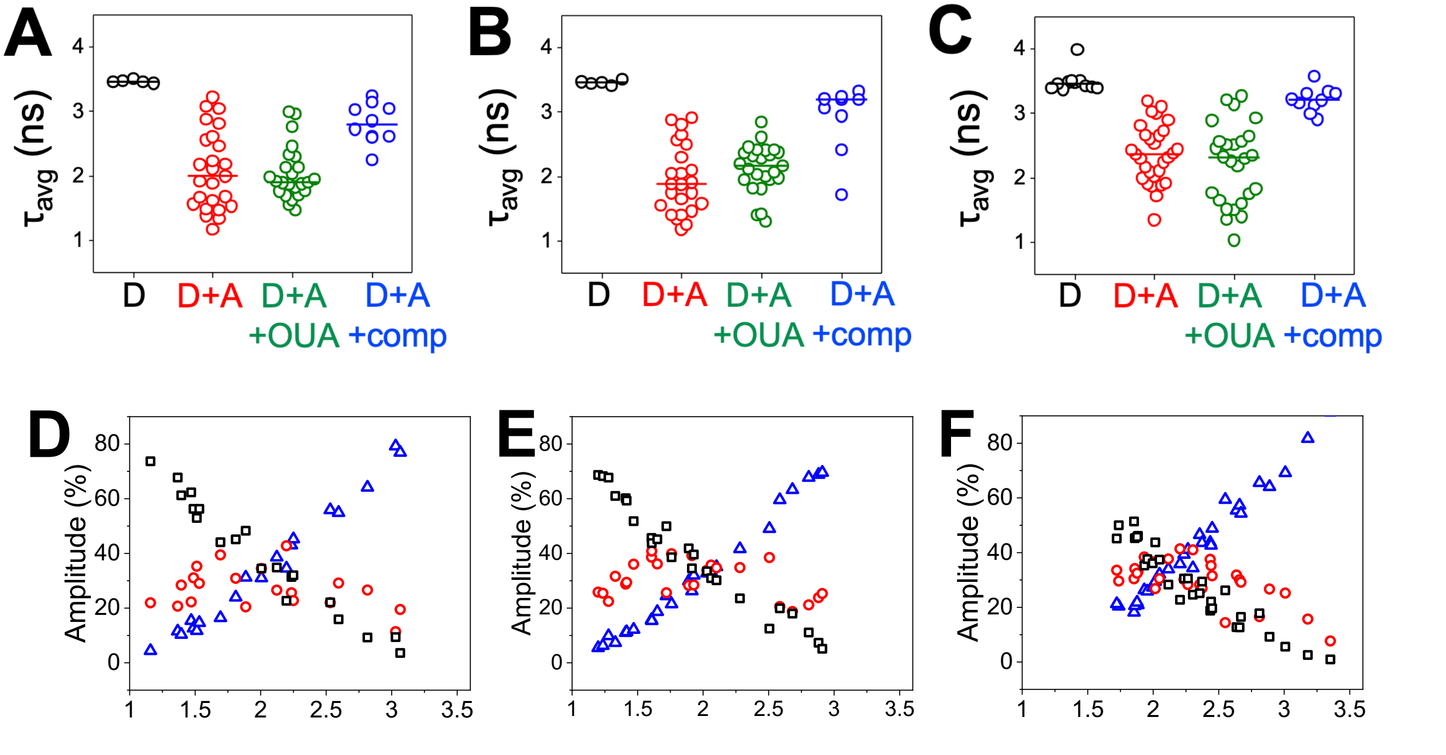


**Sup. Fig. 2**: Quantification of fluorescence decays. (**A**) A decreased fluorescence lifetime indicates FRET between donor-labeled $\alpha_1$ (D) and acceptor labeled PLM (A). FRET was not affected by addition of ouabain (OUA). FRET was reduced by coexpression of unlabeled PLM. (**B**) As in (A), but with the $\alpha_2$ isoform. (**C**) As in (A), $\alpha_3$ isoform. (**D**) The relative population of FRET species for $\alpha_1$. (**E**) The relative population of FRET species for $\alpha_2$. (**F**) The relative population of FRET species for $\alpha_3$. We observed no differences for α subunit isoforms, so the data were combined and analyzed collectively.

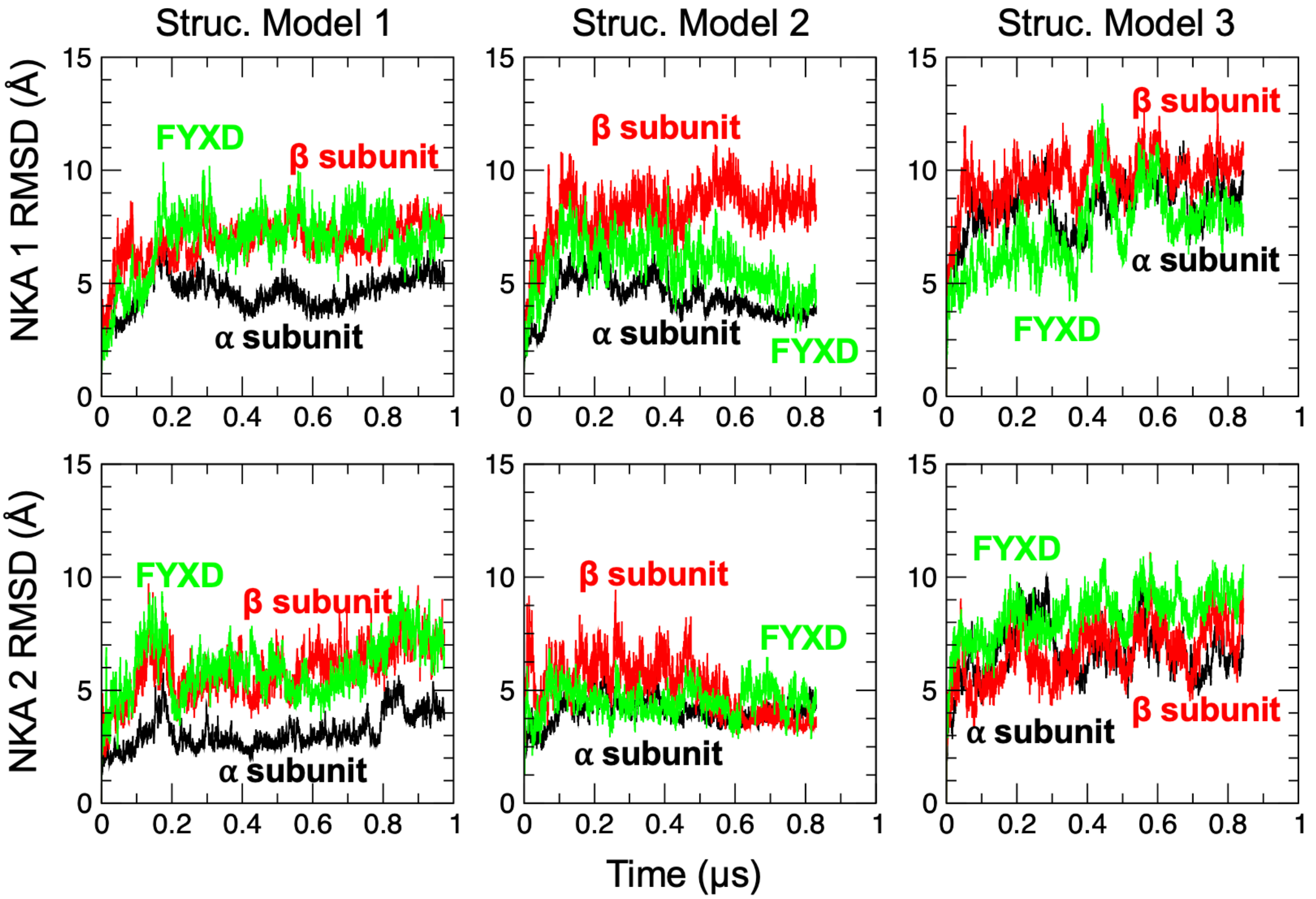


**Sup. Fig. 3**: RMSD for α (black), β (red), and FYXD (green) was calculated by aligning the backbone of the entire dimer with the structure at the beginning of each MD trajectory.

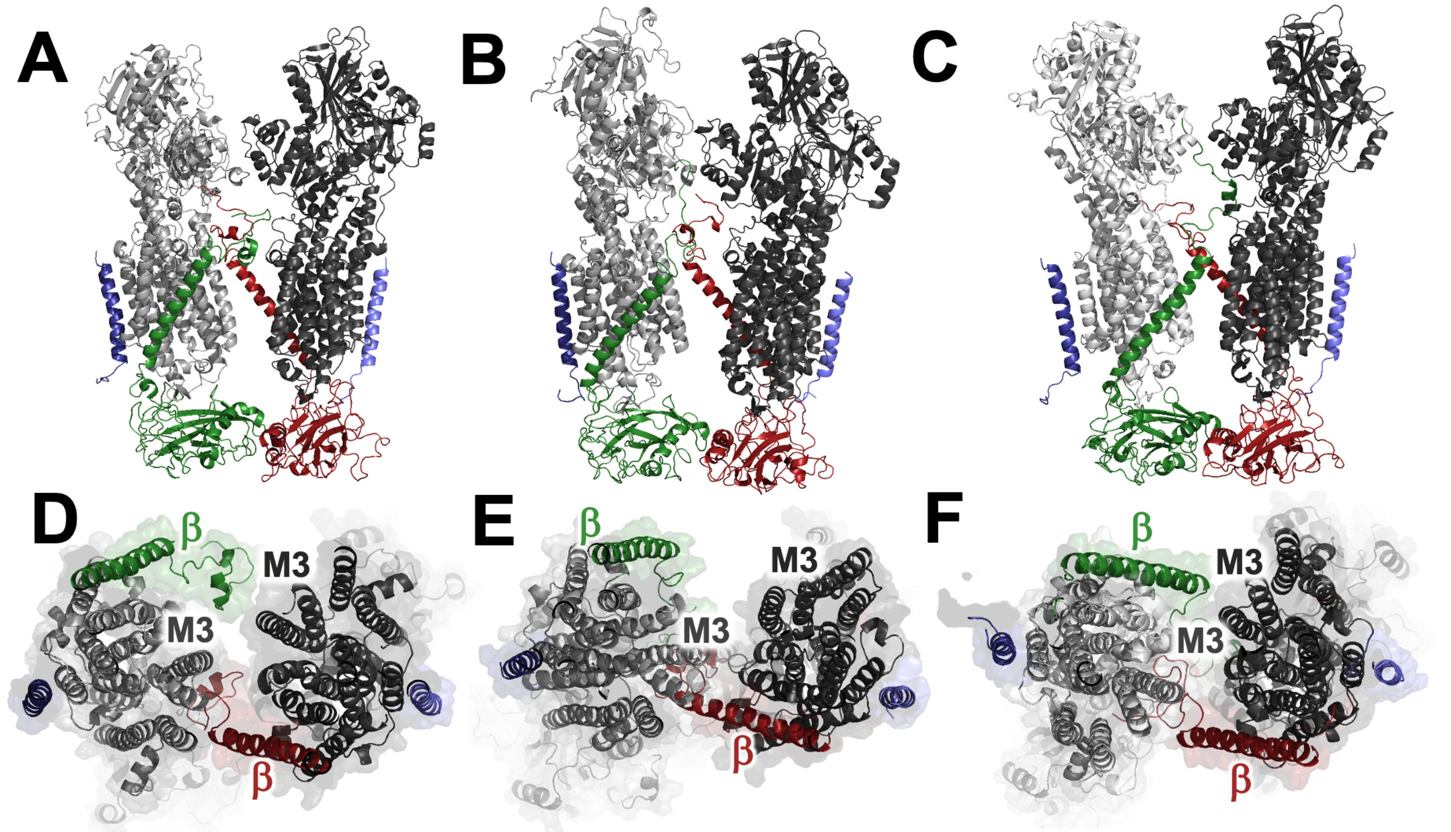


**Sup. Fig. 4**: Structures of alternative Structural Models 1, 2, and 3, showing α subunits in gray, β subunits in red and green, and FXYD proteins in blue or lavender.

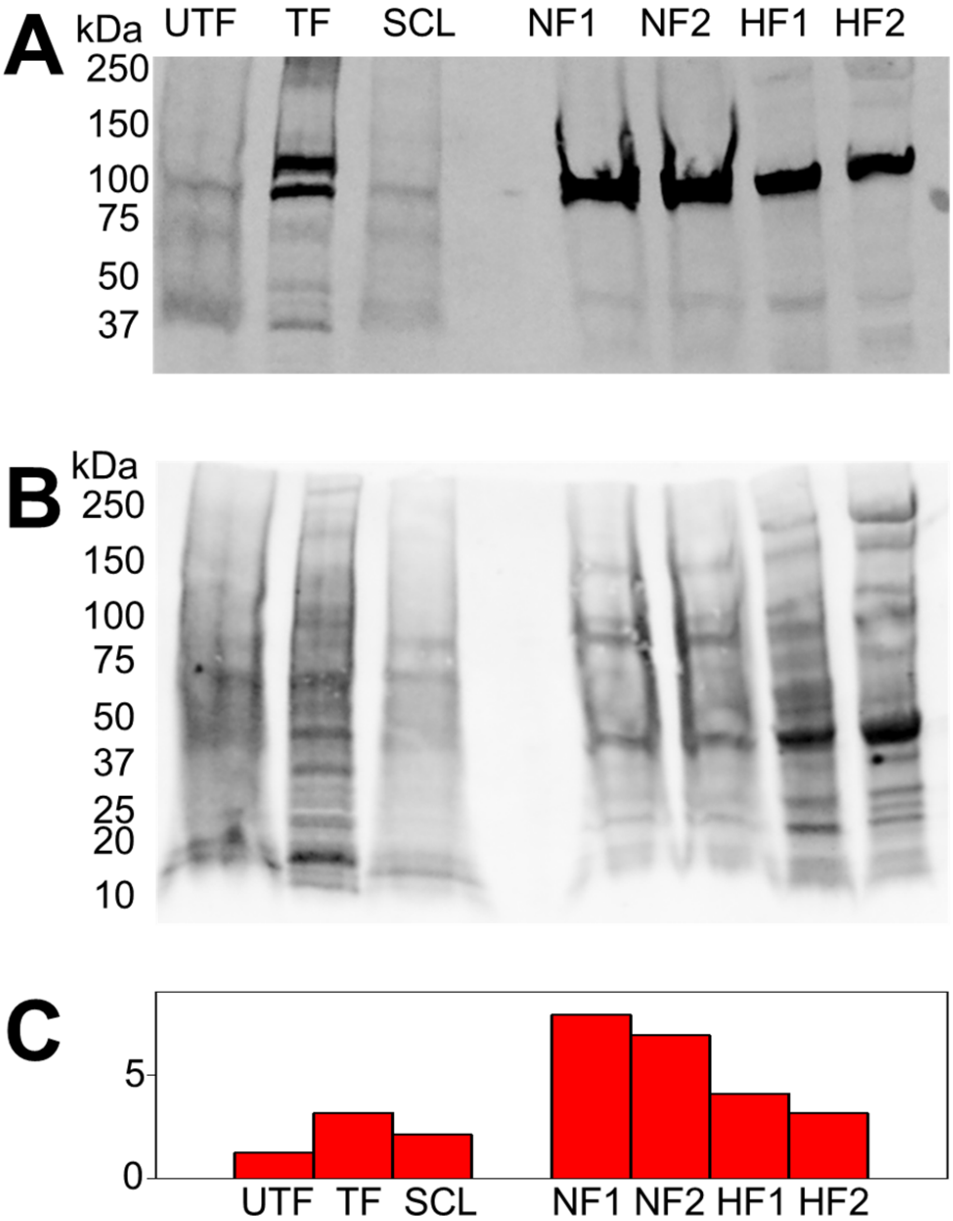


**Sup. Fig. 5**: Western blot analysis showed minimal endogenous NKA expression in un-transfected HEK cell microsomal fractions (UTF). (**A**) Increased NKA $\alpha_1$ expression was observed in microsomes from cells transfected with human mCer- $\alpha_1$ NKA (TF) and in microsomes isolated from a rat mCer-$\alpha_1$ NKA stable cell line (SCL). Note that the mobilities of microsomal fractions prepared from fluorescently labeled constructs overexpressed in HEK cells are shifted by approximately 30 kDa compared to endogenous NKA. NKA was highly expressed in microsomal fractions isolated from myocardium of non-failing human hearts (NF1-2). We observed decreased expression of NKA in failing human heart (HF1-2) compared to non-failing hearts. (**B**) Total protein staining using Revert solution. (**C**) Quantification of NKA expression in HEK cells and human non-failing and failing hearts. The signal of NKA antibody was normalized to total protein content.
